# Sequences of Intonation Units form a ~1 Hz rhythm

**DOI:** 10.1101/765016

**Authors:** Maya Inbar, Eitan Grossman, Ayelet N. Landau

## Abstract

Studies of speech processing investigate the relationship between temporal structure in speech stimuli and neural activity. Despite clear evidence that the brain tracks speech at low frequencies (~1 Hz), it is not well understood what linguistic information gives rise to this rhythm. Here, we harness linguistic theory to draw attention to Intonation Units (IUs), a fundamental prosodic unit of human language, and characterize their temporal structure as captured in the speech envelope, an acoustic representation relevant to the neural processing of speech.

IUs are defined by a specific pattern of syllable delivery, together with resets in pitch and articulatory force. Linguistic studies of spontaneous speech indicate that this prosodic segmentation paces new information in language use across diverse languages. Therefore, IUs provide a universal structural cue for the cognitive dynamics of speech production and comprehension.

We study the relation between IUs and periodicities in the speech envelope, applying methods from investigations of neural synchronization. Our sample includes recordings from every-day speech contexts of over 100 speakers and six languages. We find that sequences of IUs form a consistent low-frequency rhythm and constitute a significant periodic cue within the speech envelope. Our findings allow to predict that IUs are utilized by the neural system when tracking speech, and the methods we introduce facilitate testing this prediction given physiological data.

## Introduction

Speech processing is commonly investigated by the measurement of brain activity as it relates to the acoustic speech stimulus^1–3^. Such research has revealed that neural activity tracks amplitude modulations present in speech. It is generally agreed that a dominant element in the neural tracking of speech is a 5 Hz rhythmic component, which corresponds to the rate of syllables in speech^4–8^. The speech stimulus is also tracked at lower frequencies (<5 Hz), e.g.^2,3^, but the functional role of these fluctuations is not fully understood. They are assumed to relate to the “musical” elements of speech which are above the word level – called prosody. However, prosody in the neuroscience literature is rarely investigated for its structure and function in cognition.

In contrast, the role of prosody in speech and cognition are extensively studied within the field of linguistics. Such research identifies prosodic segmentation cues that are common to all languages, and that characterize what is termed Intonation Units (IUs; Figure 1a)^9–11^. Importantly, in addition to providing a systematic segmentation to ongoing naturalistic speech, IUs capture the pacing of information, parceling a maximum of one new idea per IU. Thus, IUs provide a valuable construct for quantifying how ongoing speech serves cognition in individual and interpersonal contexts. The first goal of our study is to introduce this understanding of prosodic segmentation from linguistic theory to the neuroscientific community. The second goal is to put forth a temporal characterization of IUs, and hence offer a precise, theoretically-motivated interpretation of the low-frequency auditory tracking and its relevance to cognition.

**Figure 1.**
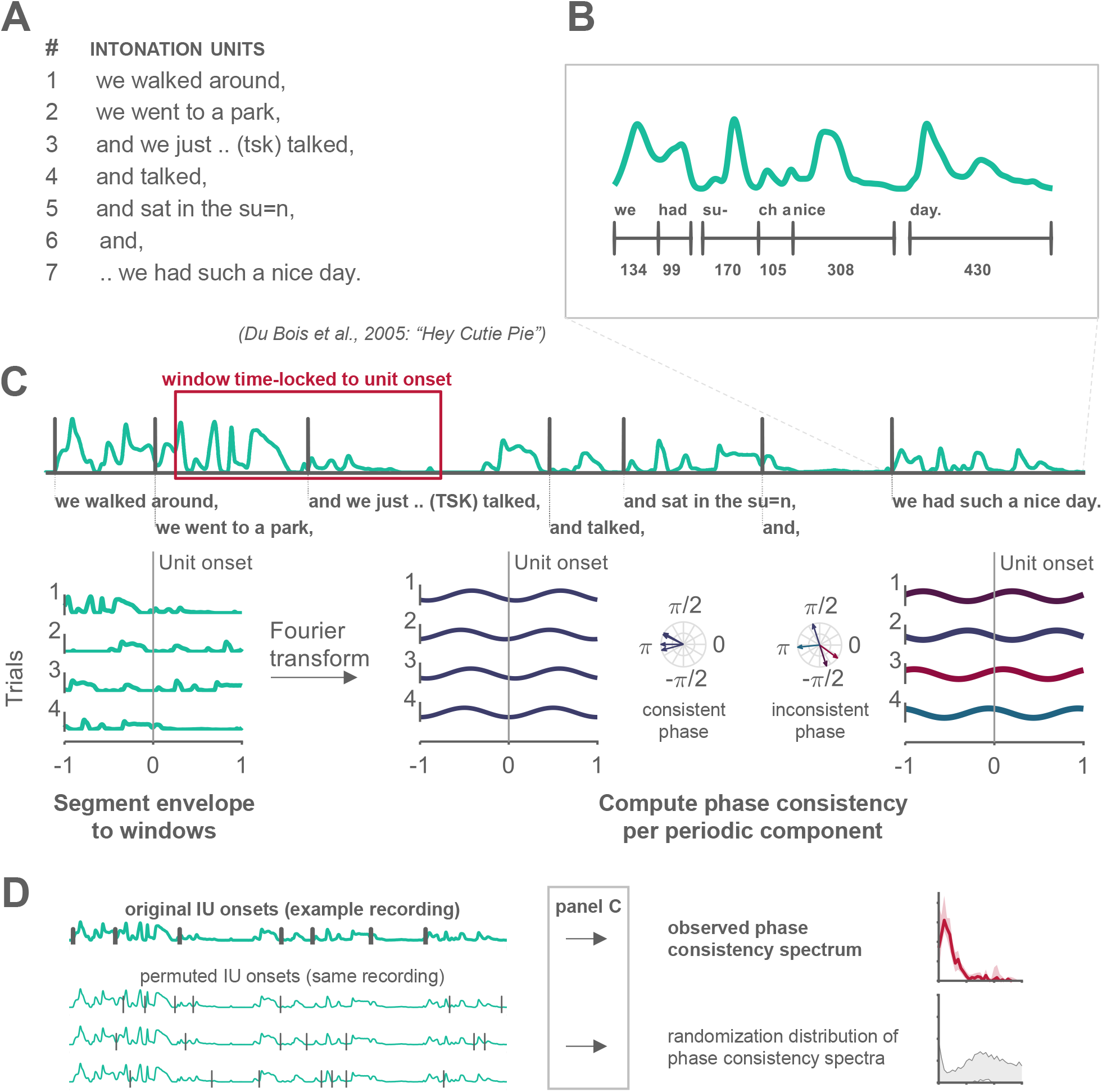
Analysis pipeline. (a) An example intonation unit sequence from a conversation in Du Bois et al.^20^. (b) Illustration of one of the characteristics contributing to the delimitation of IUs: the fast-slow dynamic of syllables. A succession of short syllables is followed by comparatively longer ones; new units are cued by the resumed rush in syllable rate following the lengthening (syllable duration measured in ms). (c) Illustration of the phase-consistency analysis: 2-second windows of the speech envelope (green) were extracted around each IU onset (gray vertical line), decomposed and compared for consistency of phase angle within each frequency. (d) Illustration of the IU-onset permutation in time, which was used to compute the randomization distribution of phase consistency spectra (see Materials and Methods).

When speakers talk, they produce their utterances in chunks with a specific prosodic profile, a profile which is attested in all human languages regardless of their phonological and morphosyntactic structure. The prosodic profile intersects rhythmic, melodic, and articulatory characteristics^9–11^. Rhythmically, chunks may be delimited by pauses, but more importantly, by a fast-slow dynamic of syllables (Figure 1b). Melodically, chunks have a continuous pitch contour, which is typically sharply reset at the onset of a new unit. In terms of articulation, the degree of contact between articulators is strongest at the onset of an IU^12^, a dynamic that generates resets in volume at the onsets of units.

Across languages, these prosodically-defined units are also functionally comparable in that they pace the flow of information in the course of speech^9,13–15^. For example, when speakers develop a narrative, they do so gradually, introducing the setting, participants and the course of events in sequences of IUs, where no more than one new piece of information relative to the preceding discourse is added per IU (Box 1). This has been demonstrated both by means of qualitative discourse analysis^9,e.g., 13,16^, and by quantifying the average amount of content items per IU. Specifically, the amount of content items per IU has been found to be very similar across languages, even when they have strikingly different grammatical profiles^10^. Another example for the common role of IUs in different languages pertains to the way speakers plan their (speech) actions. When speakers coordinate a transition during a turn-taking sequence, they rely on prosodic segmentation (i.e., IUs): points of semantic/syntactic phrase closure are not a sufficient cue for predicting when a transition will take place, and IU design is found to serve a crucial role in timing the next turn-taking transition^17–19^.

### Box 1. Information is temporally structured in social interaction.

Time is crucial for organizing information in interaction, not only via prosodic cues but also through body conduct, such as gaze-direction, head and hand gestures, leg movements and body torques^52^. This is especially evident in task-oriented interaction, for example, in direction-giving sequences during navigation, or instruction sequences more broadly, where the many deictic words (e.g., this, there) can only be interpreted correctly when accompanied by a timely gesture. Following is such a fragment from a judo-instruction class^20^. The prosodic-based segmentation into IUs is represented by a break in line, such that each line corresponds to on IU. To facilitate reading, transcription conventions were simplified (the original transcription can be retrieved with the sound file from the linked corpus). Speaker overlap is marked by square brackets ([]), minimally audible to medium pauses (up to 0.6 seconds) are marked by sequences of dots (…) and punctuation marks represent different functional classes of pitch movements, indicating roughly the degree of continuity between one unit and the next (comma – continuing; period – final; double dash – cut short).

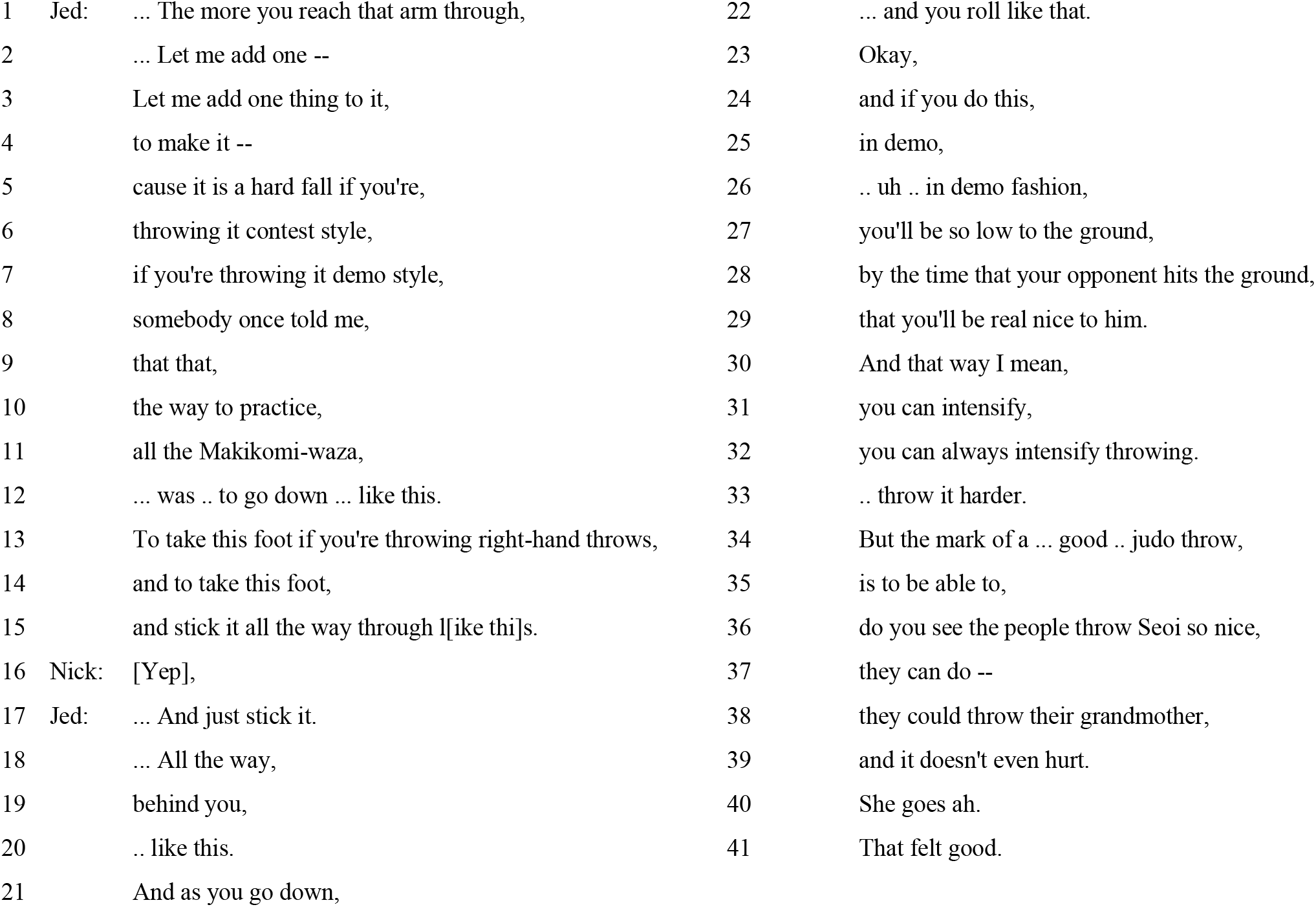
“Throw me” (SBC057: 24:19–25:19)^20^.

Here we use recordings of spontaneous speech in natural settings to characterize the temporal structure of sequences of IUs in six languages. The sample includes well-studied languages from the Eurasian macro-area, as well as much lesser-known and -studied languages, spoken in the Indonesian-governed part of Papua by smaller speech communities. Importantly, our results generalize across this linguistic diversity, despite the substantial differences in socio-cultural settings and all aspects of grammar, including other prosodic characteristics. In contrast to previous research, we estimate the temporal structure of IUs using direct time measurements rather than word or syllable counts (Box 2). In addition, we quantify the temporal structure of IUs in relation to the speech envelope, which is an acoustic representation relevant to neural processing of speech. We find that sequences of IUs form a consistent low-frequency rhythm at ~1 Hz in the six sample languages, and relate this finding to recent neuroscientific accounts of the roles of slow rhythms in speech processing.

### Box 2. What is a word?

The notion of a word has been argued to be untenable for both language-specific and cross-linguistic analyses (e.g.,^53^). We demonstrate why this is so for cross-linguistic comparison with the following example in Seneca, a member of the Northern Iroquoian branch of the Iroquoian language family spoken in Northeast America^54^.

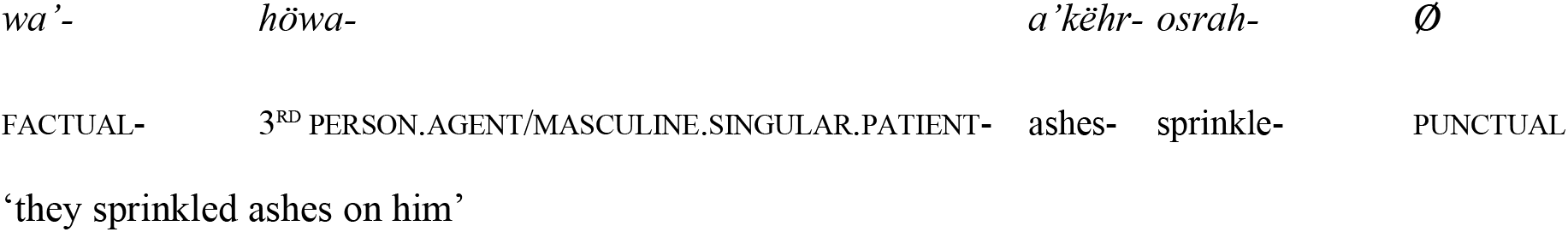
wáö́wö’géosäh.

The first line includes a word in the language, that is, a unit of meaning whose unit-ness is defined by morphosyntactic processes in the language. The second line includes a breakdown to meaning components (separated by hyphens and tabs), obtained through linguistic analysis and comparative evidence from related languages. Due to extensive sound changes over the years, Seneca shows a high degree of fusion between meaning components, that is, the boundaries between them are obscured and not necessarily available to speakers. Note also that these meaning components cannot normally appear as independent words, that is, without the neighboring meaning components. The third line includes a gloss per meaning component, differentiating between those with grammatical meaning (part of a grammatical paradigm; in small caps) and content items. The fourth line includes the corresponding English translation.

As evident from this example, a noteworthy distinction between Seneca and some better-known languages of the world such as English is that Seneca regularly packages an event, its participants and other meaning components within a single morphosyntactic word. Consequently, for one and the same message, Seneca IUs would contain fewer words compared to English IUs.

Many other linguistic constructs vary greatly from language to language. In fact, it seems that linguistic diversity is the rule rather than the exception, and that care should be taken to avoid *a priori* taxonomies that would fail when considering the next language^55,56^.

## Materials and Methods

### Data

We studied the temporal structure of IUs using six corpora of conversations and narratives that were transcribed and segmented into IUs according to the unified criteria devised by Chafe, Du Bois and colleagues^9,11^. Three of the corpora were segmented by specialist teams working on their native language: the Santa Barbara Corpus of Spoken American English^20^, the Haifa Corpus of Spoken Hebrew^21^, and the Russian Multichannel Discourse corpus^22^. The other three corpora were segmented by teams with varying degrees of familiarity with the languages, as part of a project studying the human ability to identify IUs in unfamiliar languages: the DoBeS Summits-PAGE Collection of Papuan Malay^23^, the DoBeS Wooi Documentation^24^, and the DoBes Yali Documentation^25^. Further information regarding the sample is found in Table 1 and Table S1. Appendix A in the Supplementary Information elaborates on the construction of the sample and the coding and processing of IUs. From all language samples, we extracted IU onset times, noting which speaker produced a given IU.

**Table 1.**
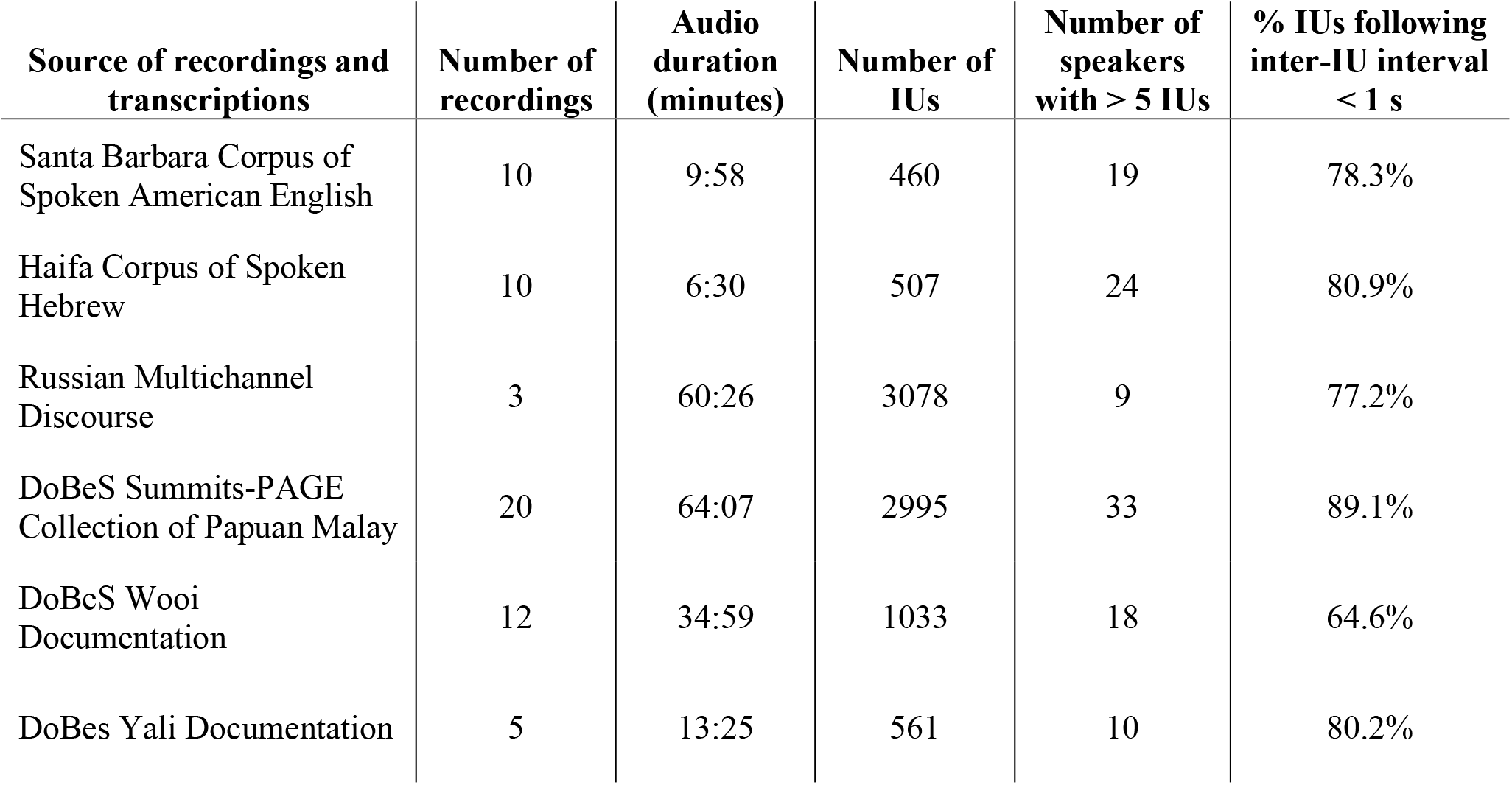
Summary information on the sample of speech segments used in the study.

| Source of recordings and transcriptions | Number of recordings | Audio duration (minutes) | Number of IUs | Number of speakers with > 5 IUs | % IUs following inter-IU interval < 1 s |
| --- | --- | --- | --- | --- | --- |
| Santa Barbara Corpus of Spoken American English | 10 | 9:58 | 460 | 19 | 78.3% |
| Haifa Corpus of Spoken Hebrew | 10 | 6:30 | 507 | 24 | 80.9% |
| Russian Multichannel Discourse | 3 | 60:26 | 3078 | 9 | 77.2% |
| DoBeS Summits-PAGE Collection of Papuan Malay | 20 | 64:07 | 2995 | 33 | 89.1% |
| DoBeS Wooi Documentation | 12 | 34:59 | 1033 | 18 | 64.6% |
| DoBes Yali Documentation | 5 | 13:25 | 561 | 10 | 80.2% |

### Phase-consistency analysis

We analyzed the relation between IU onsets and the speech envelope using a point-field synchronization measure, adopted from the study of rhythmic synchronization of neural spiking activity and Local Field Potentials^26^. In this analysis, the rhythmicity of IU sequences is measured through the phase consistency of IU onsets with respect to the periodic components of the speech envelope (Figure 1c). The speech envelope is a representation of speech that captures amplitude fluctuations in the acoustic speech signal. The envelope is most commonly understood to reflect the succession of syllables at ~5 Hz, and indeed, it includes strong 2-7 Hz modulations^4,6–8^. The vocal nuclei of syllables are the main source of envelope peaks, while syllable boundaries are the main source of envelope troughs (Figure 1b). IU onsets can be expected to coincide with troughs in the envelope, since each IU onset is necessarily also a syllable boundary. Therefore, one can expect a high phase consistency between IU onsets and the frequency component of the speech envelope corresponding to the rhythm of syllables, at ~5 Hz.

A less trivial finding would be a high phase consistency between IU onsets and other periodic components in the speech envelope. Specifically, since IUs typically include more than one syllable, such an effect would pertain to frequency components below ~5 Hz. In this analysis we hypothesized that the syllable organization within IUs gives rise to slow periodic components in the speech envelope. If low-frequency components are negligible in the speech envelope, estimating the phase of the low-frequency components at the time of IU onsets would lead to random phase angles, a result that would translate to low phase consistency (i.e., uniformity). In another scenario, if the speech envelope captures slow rhythmicity in language other than that arising from IUs, different IUs would occur in different phases of the lower frequency components, translating again to low phase consistency. In contrast to these scenarios, finding phase consistency at a degree higher than expected under the null hypothesis would indicate both that the speech envelope captures the rhythmic characteristics of IUs and would characterize the period of this rhythmicity.

We computed the speech envelope for each sound file following standard procedure^3, Figure 1c and Appendix A,8^. We extracted 2-second windows of the speech envelope centered on each IU onset, and decomposed them using Fast Fourier Transform (FFT) with a single Hann window, no padding, following demeaning. This yielded phase estimations for frequency components at a resolution of 0.5 Hz. We then measured the consistency in phase of each FFT frequency component across speech segments using the pairwise-phase consistency metric (PPC)^26^, yielding a consistency spectrum. We calculated consistency spectra separately for each speaker that produced > 5 IUs and averaged the spectra within each language. Note, that the PPC measure is unbiased by the number of 2-second envelope windows entering the analysis^26^, and likewise that in a turn-taking sequence, it is inevitable that part of the 2-second envelope windows capture speech by more than one participant. We also conducted the analysis using 4-second windows of the speech envelope, allowing for a 0.25 Hz resolution but at the expense of less data entering the analysis. Further information regarding this additional analysis can be found in Appendix B in the Supplementary Information.

### Statistical assessment

We assessed the statistical significance of peaks in the average consistency spectra using a randomization procedure (Figure 1d). Per language, we created a randomization distribution of consistency estimates with 1000 sets of average surrogate spectra. These surrogate spectra were calculated using the speech envelope as before, but with temporally permuted IU onsets that maintained the association with envelope troughs. Troughs are defined by a minimum magnitude of 0.01 (on a scale of 0-1), and with a minimal duration between troughs of 200 ms, as would be expected from syllables, on average. By constraining the temporal permutation of IU onsets, we address the fact that each IU onset is necessarily a syllable onset, and therefore is expected to align with a trough in the envelope. We then calculated, for each frequency, the proportion of consistency estimates (in the 1000 surrogate spectra) that were greater than the consistency estimate obtained for the observed IU sequences. We corrected p-values for multiple comparisons across frequency bins ensuring that on average, False Discovery Rate (FDR) will not exceed 1%^27,28^.

## Results

We studied the temporal structure of IU sequences through their alignment with the periodic components of the speech envelope, using a phase-consistency analysis. This analysis builds on a cognitively-oriented linguistic theory which is supported by empirical speech analysis with wide cross-linguistic validity. We hypothesized that one of the characteristics of IUs – the fast-slow dynamic of syllables – would give rise to slow periodic modulations in the speech envelope. Figure 2 displays the observed phase consistency spectra in the six sample languages. IU onsets appear at significantly consistent phases of the low-frequency components of the speech envelope, indicating that their rhythm is captured in the speech envelope, hierarchically above the syllabic rhythm at ~5 Hz (English: 0.5-1.5 Hz; Hebrew: 1-1.5, 2.5-3 Hz; Russian: 0.5-3 Hz; Papuan Malay: 0.5-3.5 Hz; Wooi: 0.5-3.5 Hz; and Yali: 0.5-4 Hz, all *p*’s < 0.001). Of note, the highest phase consistency is measured at 1 Hz in all languages except Hebrew, in which the peak is at the neighboring frequency bin, 1.5 Hz.

**Figure 2.**
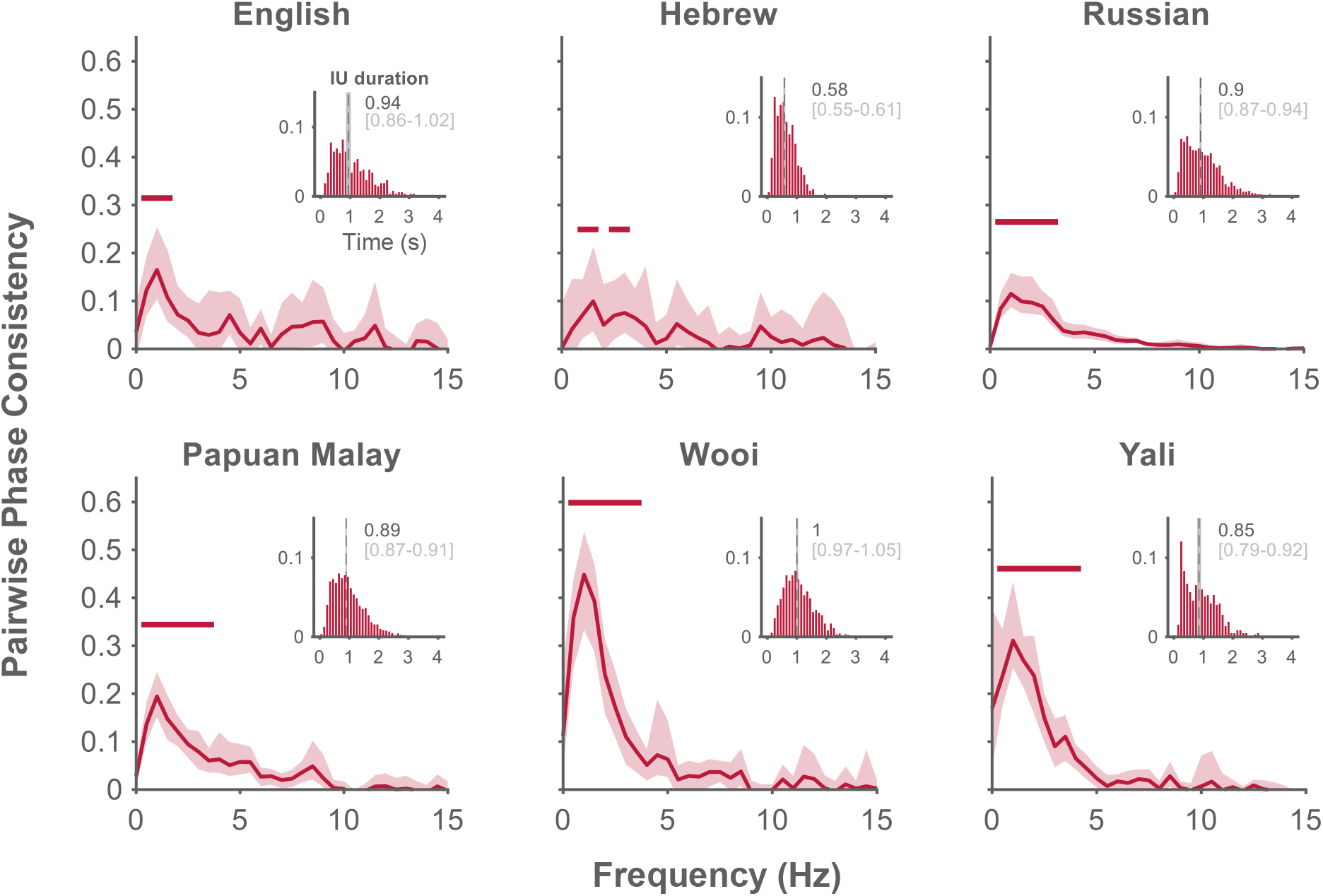
Characterization of the temporal structure of Intonation Units. Phase-consistency analysis results include the average of phase consistency spectra across speakers for each language. Shaded regions denote bootstrapped 95% confidence intervals^32^ of the averages. Significance is denoted by a horizontal line above the spectra, after correction for multiple comparisons across neighboring frequency bins using an FDR procedure. Inset: Probability distribution of IU durations within each language corpus, calculated for 50 ms bins and pooled across speakers. Overlaid are the medians (dashed line; dark gray) and the bootstrapped 95% confidence intervals of the medians (light gray).

To complement the results of the phase-consistency analysis, we estimated the median duration of IUs (Figure 2, insets). The bootstrapped 95% confidence intervals of this estimate are mostly overlapping, to a resolution of 0.1 s, for all languages but Hebrew. For Hebrew, the median estimate indicates a shorter IU duration, which may underlie a faster rhythm of IU sequences. Note, however, that duration is only a proxy for the rhythmicity of IU sequences, as IUs do not always succeed each other without pause (Figure S2, insets). We find reassuring the consistent trends in the two analyses, but do not pursue the post-hoc hypothesis that Hebrew deviates from the other languages. In the planned phase consistency analysis, the range of significant frequency components is consistent across languages.

We sought to confirm that this effect was not a result of an amplitude transient at the beginning of IU sequences. To this end, we repeated the analysis, submitting only IUs that followed an inter-IU interval below 1 s, that is, between 65%-89% of the data, depending on the language (Table 1). The consistency estimates at 1 Hz were still larger than expected under the null hypothesis that IUs lack a definite rhythmic structure (Figure S2).

Our results are consistent with preliminary characterizations of the temporal structure of IUs^13,29,30^. The direct time measurements we used obviate the pitfalls of length measurements in word count or syllable count, (e.g.,^9,10,14,30^). The temporal structure of IUs cannot be inferred from reported word counts, because what constitutes a word varies greatly across languages (Box 2). Syllable count per IU may provide an indirect estimation of IU length, especially if variation in syllable duration is taken into account (e.g.,^31^), but it does not capture information about the temporal structure of IU *sequences*.

## Discussion

Neuroscientific studies suggest that neural oscillations participate in segmenting the auditory signal and encoding linguistic units during speech perception. Many studies focus on the role of oscillations in the theta range (~5 Hz) and in the gamma range (>40 Hz). The levels of segmentation attributed to these ranges are the syllable level and the fine-grain encoding of phonetic detail, respectively^1^. Studies identify also slower oscillations, in the delta range (~1 Hz), and have attributed them to segmentation at the level of phrases, both prosodic and semantic/syntactic^2,3,33–37^. Previous studies have consistently demonstrated a decrease in high-frequency neural activity at points of semantic/syntactic completion^38^ or natural pauses between phrases^39^. This pattern of activity yields a slow modulation aligned to phrase structure. We harness linguistic theory to offer a conceptual framework for such slow modulations. We quantify the phase consistency between acoustical slow modulations and IU onsets, and demonstrate for the first time that prosodic units with established functions in cognition give rise to a low-frequency rhythm in the auditory signal available to listeners.

Previous research has proposed to dissociate delta activity that represents acoustically-driven segmentation following prosodic phrases from delta activity that represents knowledge-based segmentation of semantic/syntactic phrases^40^. From the perspective of studying the temporal structure of spontaneous speech, we suggest that the distinction maintained between semantic/syntactic and prosodic phrasing might be superficial. That is because the semantic/syntactic building blocks always appear within prosodic phrases in natural language use^41–44^. Studies investigating semantic/syntactic building blocks often compare the temporal dynamics of intact grammatical structure to word lists or grammatical structure in an unfamiliar language (e.g.,^33,36,38^). We argue that such studies need to incorporate the possibility that ongoing processing dynamics might reflect perceptual chunking, owing to the ubiquity of prosodic segmentation cues in natural language experience. This possibility is further supported by the fact that theoretically-defined semantic/syntactic boundaries are known to enhance the perception of prosodic boundaries, even when those are artificially removed from the speech segment. In a study that investigated the role of syntactic structure in guiding the perception of prosody in naturalistic speech^45^, syntactic structure was found to make an independent contribution to the perception of prosodic grouping. Another study equated prosodic boundary strength experimentally (controlling in a parametric fashion word duration, pitch contour, and following-pause duration), and found the same result: semantic/syntactic completion contributed to boundary perception^46^. Even studies that use visual serial word presentation paradigms rather than auditory stimuli are not immune to an interpretation of prosodically-guided perceptual chunking, which is known to affect silent reading^47^ (for a review see^48^).

Independently of whether delta activity in the brain of the listener represents acoustic landmarks, abstract knowledge, or the prosodically-mediated embodiment of abstract knowledge^42^, our results point to another putative role for slow rhythmic brain activity. We find that orthogonal to different grammatical systems, speakers and speech modes, speakers express their developing ideas at a rate of approximately 1 Hz. Previous studies have shown that in the brains of listeners, a wide network interacts with low-frequency auditory-tracking activity, suggesting an interface of prediction and attention-related processes, memory and the language system^2,37,49–51^. We expect that via such low-frequency interactions, this same network constraints spontaneous speech production, orchestrating the management and communication of conceptual foci^9^.

Finally, our findings render plausible several hypotheses within the field of linguistics. At a basic level, the consistent duration of IUs may provide a temporal upper bound to the construal of other linguistic units (e.g., morphosyntactic words).

## Acknowledgement

MI is supported by the Humanities Fund PhD program in Linguistics and the Jack, Joseph and Morton Mandel School for Advanced Studies in the Humanities. Our work would not have been possible without the substantial efforts carried by the creators of the corpora, their teams, and the people they recorded all over the world.

## Author contributions

This work is an outcome of author MI’s MA thesis under the joint supervision of authors ANL and EG. The authors were jointly active in all stages of this research and its publication.

## Data availability statement

The custom-written code producing the analyses and figures will be made available through the Open Science Framework repository upon publication. As for data, IU time stamps for Hebrew and English can be retrieved from the authors upon request. IU time stamps for the rest of the languages as well as all audio files can be retrieved from the cited corpora.

