## Supplementary Information for "Sequences of Intonation Units form a ~1 Hz rhythm"

**This Supplementary Information file includes:**

Appendix A: Supplementary Materials and Methods

Supplementary Table S1

Appendix B: Supplementary Analyses

Supplementary Figures S1-S2

References for Supplementary Information citations

### Appendix A: Supplementary Materials and Methods

*Data*

We analyzed data from six corpora of spontaneous speech, which fall into two groups. The first group consists of corpora in English, Hebrew and Russian, transcribed and segmented by professional teams, each working on their native language: the Santa Barbara Corpus of Spoken American English[1], the Haifa Corpus of Spoken Hebrew[2], and the Russian Multichannel Discourse corpus[3]. We constructed a sample from each of the three corpora according to the following protocol: we sampled randomly 10 recordings from the English and Hebrew corpora, and included all 9 recordings available as a preview to the Russian corpus. The English and Hebrew corpora include transcriptions segmented into intonation units (IUs) that are not timestamped to the audio file. Author MI manually measured the IU- onset and -offset times in the first 30 seconds of each recording, continuing until the next speaker change in each recording. Pauses between IUs were not considered as part of the neighboring IUs' duration, in deviation from the transcription guidelines of Du Bois et al.[4]. Spectrogram analyses in Praat 6.0.23[5] accompanied this procedure, including pitch and intensity contours produced by Praat's default settings. Measurements in milliseconds were entered into ELAN 4.9.4[6], on separate tiers for different speakers. The Russian corpus includes the onset and offset times of IUs throughout each recording, measured in a similar fashion. Finally, in English and Hebrew, additional information was recorded for each IU, but was not further analyzed: the intonation contour attributed by the corpus constructors, the position within the turn, and a categorization to type of IU – Fragmentary, Regulatory or Substantive[7]. We replicated the results of the main analysis with 20 recordings from a Russian corpus that included narratives recorded by individual speakers rather than multi-speaker conversations[8]. In that dataset, although the sample of speakers was larger, the total recording time was less than a quarter of the length of the dataset analyzed in the current report. Nevertheless, the results were identical and we do not report them here.

The second group consists of corpora in Papuan Malay, Wooi and Yali: the DoBeS Summits-PAGE Collection of Papuan Malay[9], the DoBeS Wooi Documentation[10], and the DoBes Yali Documentation[11]. Papuan Malay (ISO 639-3 code: pmy) is the *lingua franca* of West Papua, the western half of the island of New Guinea governed by Indonesia. Wooi (ISO 639-3 code: wbw) and Yali (ISO 639-3 code: yac) belong to different language families, have very different grammatical profiles, and are spoken in different regions in West Papua – on the coast and in the highlands, respectively. All three corpora include retellings of the plot of a silent film by a narrator to an interlocuter. More data on these languages and corpora can be found in Himmelmann et al.[12], a study that was dedicated to the human ability to identify IUs in unfamiliar languages. In this study, researchers with varying degrees of familiarity with Papuan Malay, Wooi and Yali listened to recordings in these languages and in their native language and segmented all the recordings into IUs. The quantified agreement between researchers as to the correct segmentation was equally high for the unfamiliar languages as it was for the familiar one. Nikolaus Himmelmann kindly provided us with the consensus transcriptions that the team constructed. These were already entered into ELAN[6] and time-aligned to the recording. We retrieved the respective audio files from the DoBeS archive. The Eastern Indonesia data in Himmelmann et al.[12] included an additional language that we did not attempt to analyze because it included only two recordings. Additionally, we could not retrieve the audiofile of one of the original 6 narratives from the Yali consensus set, so only 5 Yali recordings were analyzed.

Data were extracted from all the ELAN files with the aid of custom-written scripts in R 3.6.0[13] to data frames including IU onset and offset times, the computed duration, and the speaker of each IU. All analyses were performed in MATLAB(MATLAB version 9.1.0.441655, 2016b) on a CentOS Linux 7 computer, using custom-written scripts and the FieldTrip toolbox[15]. Figures were produced using MATLAB(MATLAB version 9.3.0.713579, 2017b) on a Windows 10 Home computer.

*Speech envelope computation*

The amplitude envelope was computed for the audio file of each speech segment following the methods used in several studies demonstrating neural entrainment to the envelope [e.g., 17], methods that partly follow Chandrasekaran et al.[18].

Speech segments that were recorded in stereo were converted to mono by averaging the channels. Recordings with a sampling rate above 20 kHz were downsampled to 20 kHz. Speech segments were band-pass filtered into 10 bands between 200 Hz and half the audio file's sampling frequency, with cut-off points designed to be equidistant on the human cochlear map. Amplitude envelopes for each band (the narrowband envelopes) were computed as absolute values of the Hilbert transform. These narrowband envelopes were downsampled to 1000 Hz and subsequently averaged, yielding the wideband envelope. The wideband envelope was smoothed using a 50 ms sliding Gaussian filter and divided by its maximal value to be on a scale of 0-1.

| Corpus | Recordings | |
|---|---|---|
| Santa Barbara Corpus of Spoken American English[1] | Just_wanna_hang<br>New_Yorkers_anonymous<br>Noise_pollution<br>Oh_you_need_a_breadbox<br>Shaggy_dog_story | Swingin_kid<br>The_mama_of_dada<br>Throw_me<br>What_is_a_brand_inspection<br>You_baked |
| Haifa Corpus of Spoken Hebrew[2] | Changing_the_education_system<br>Conversation_about_conversation_and_babies<br>Costume_for_purim<br>Grenade<br>Gym | Korin_is_in_love<br>Lack_of_tact_and_sensitivity<br>Looking_for_work<br>Luck_in_cards_and_girls<br>Parenthood_and_education |
| Russian Multichannel Discourse corpus[3] | Pears04<br>Pears23 | Pears22 |
| DoBeS Summits-PAGE Collection of Papuan Malay[9] | PMY_pear_Boas_KONS<br>PMY_pear_Carl_KONS<br>PMY_pear_Fant_KONS<br>PMY_pear_Irma_KONS<br>PMY_pear_Lala_KONS<br>PMY_pear_Laod_KONS<br>PMY_pear_Lia_KONS<br>PMY_pear_Lisa_KONS<br>PMY_pear_Maik_KONS<br>PMY_pear_Maya_KONS | PMY_pear_Miry_KONS<br>PMY_pear_Moha_KONS<br>PMY_pear_Nova_KONS<br>PMY_pear_Sofi_KONS<br>PMY_pear_Suar_KONS<br>PMY_pear_Titi_KONS<br>PMY_pear_Wolt_KONS<br>PMY_pear_Yanu_KONS<br>PMY_pear_Yul1_KONS<br>PMY_pear_Yul2_KONS |
| DoBeS Wooi Documentation[10] | WBW_pear_Abra_KONS<br>WBW_pear_Agus_KONS<br>WBW_pear_Davi_KONS<br>WBW_pear_Ello_KONS<br>WBW_pear_Feli_KONS<br>WBW_pear_Heri_KONS | WBW_pear_John_KONS<br>WBW_pear_Kosm_KONS<br>WBW_pear_Mart_KONS<br>WBW_pear_Oni_KONS<br>WBW_pear_Sofi_KONS<br>WBW_pear_Yuli_KONS |
| Yali Documentation[11] | YAC_pear_Edis_KONS<br>YAC_pear_Edo_KONS<br>YAC_pear_Ibra_KONS | YAC_pear_Yust_KONS<br>YAC_pear_Yusu_KONS |

*Table S1.* List of recordings used in the study.

### Appendix B: Supplementary Analyses

*Larger windows for spectral decomposition in the phase consistency analysis*

In the main analysis, we extracted 2-second windows of the speech envelope centered on each IU, decomposed them and submitted the resulting phase estimations to a consistency analysis. The inclusion of as much data as possible guided this choice, as the size of the window influences the amount of data submitted to the analysis. With 2-second windows centered on IU onsets, only the first and last seconds of each recording are excluded from the analysis. This corresponded to 5% of the data in our smallest dataset, the Hebrew dataset, whose recordings were also generally shorter. In comparison, 4-second windows would lead to the exclusion of the first and last two seconds of each recording, corresponding to 12% of the Hebrew data. However, since larger analysis windows allow for phase estimations in a finer frequency resolution and based on more cycles, we repeated the analysis using 4-second windows of the speech envelope around IU onsets (Figure S1).

IU onsets still appear at significantly consistent phases of the low-frequency components of the speech envelope. The significant ranges using the same statistical criteria (i.e. while still ensuring that FDR < 1%) now exclude the higher frequencies (English: 1-1.25 Hz; Hebrew: 1 Hz; Russian: 0.5-3.25 Hz; Papuan Malay: 0.5-3.25 Hz; Wooi: 0.25-2.5 Hz; and Yali: 0.5-2.75 Hz, all $p$'s $< 0.001$). The highest phase consistency is measured at 1 Hz for English and Hebrew, at the neighboring frequency bin 0.75 Hz for Papuan Malay, Russian and Wooi, and at 0.5 Hz for Yali.

The insets in Figure S1 present, for each language, the grand average of the speech envelope segments that were submitted to spectral decomposition. Each single envelope window was demeaned, the windows around IUs produced by the same speaker were averaged, and the grand average was then calculated across all the within-speaker average time courses. The grand average time courses clearly show that the envelope captures IU onsets. Although it is the individual segments that are submitted to the spectral decomposition and not the average of segments, this representation allows to appreciate the invariant component across the windows. This component gives rise to the measured phase consistency.

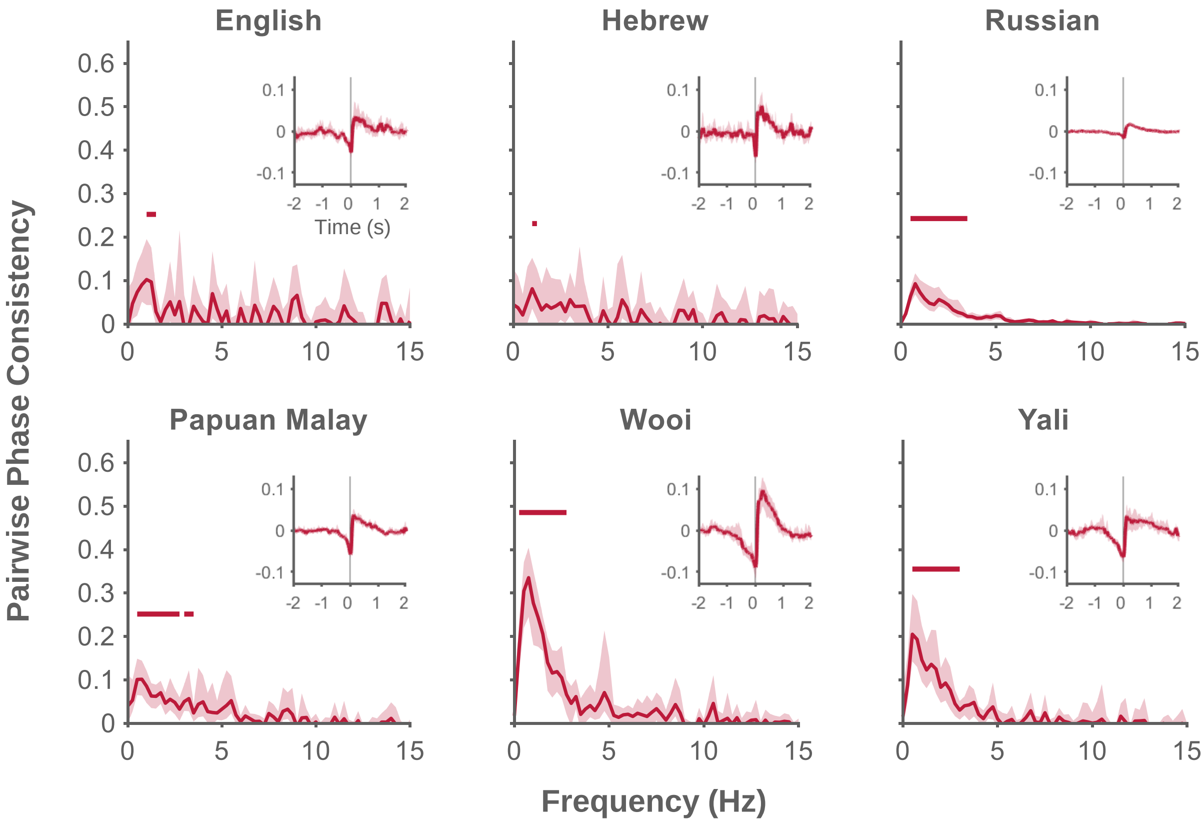


**Figure S1. Larger analysis windows do not qualitatively change the characterization of the temporal structure of Intonation Units** (item corresponding to Figure 2). Phase-consistency analysis results include the average of phase consistency spectra across speakers for each language. Shaded regions denote bootstrapped 95% confidence intervals of the averages. Significance is denoted by a horizontal line above the spectra, after correction for multiple comparisons across neighboring frequency bins using an FDR procedure. Inset: Average envelope segments time-locked to IU onsets (depicted by vertical gray line). Shaded regions denote bootstrapped 95% confidence intervals of the averages.

*Addressing a question of transient effects*

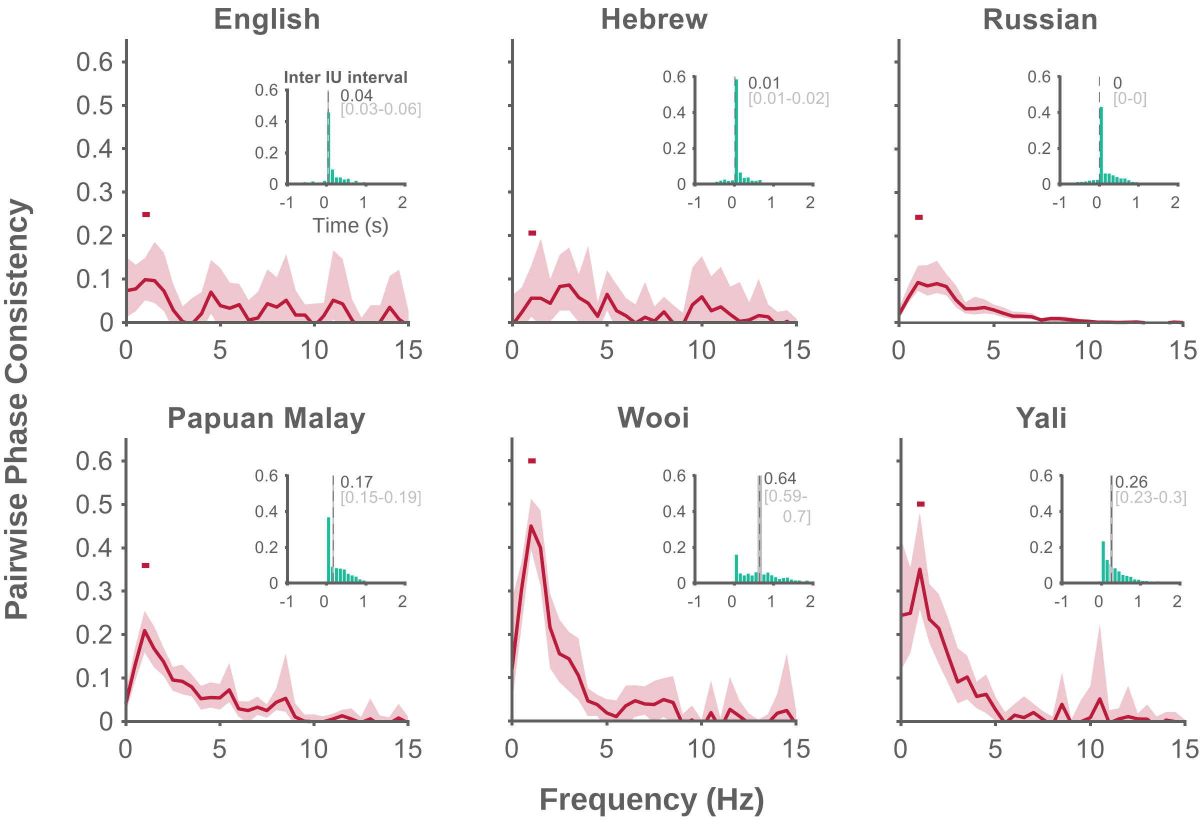


**Figure S2. Removal of IUs following long pauses does not qualitatively change the characterization of the temporal structure of Intonation Units** (item corresponding to Figure 2). Phase-consistency analysis results include the average of phase consistency spectra across speakers for each language. Shaded regions denote bootstrapped 95% confidence intervals of the averages. Significance was assessed for the 1 Hz frequency component of each spectrum and is denoted by a horizontal line above the peak. Inset: Probability distribution of all inter-IU-interval durations within each language corpus, calculated for 50 ms bins and pooled across speakers. Overlaid are the medians (dashed line; dark gray) and the bootstrapped 95% confidence intervals of the medians (light gray).
